# *Mafb* Deficiency in Myeloid Cells Increases Susceptibility to *Mycobacterium tuberculosis* Infection in Mice

**DOI:** 10.1101/2025.07.06.663407

**Authors:** Haruka Hikichi, Hajime Nakamura, Omori Shiho, Shintaro Seto, Minako Hijikata, Michito Hamada, Satoru Takahashi, Naoto Keicho

**Affiliations:** Department of Pathophysiology and Host Defense, The Research Institute of Tuberculosis, Japan Anti-Tuberculosis Association, Tokyo, Japan; Department of Basic Mycobacteriosis, Nagasaki University Graduate School of Biomedical Sciences, Nagasaki, Japan; Laboratory Animal Resource Center in Transborder Medical Research Center, and Department of Anatomy and Embryology, Institute of Medicine, University of Tsukuba, Ibaraki, Japan; Center for Medical Sciences, Ibaraki Prefectural University of Health Sciences, Ibaraki, Japan; The Research Institute of Tuberculosis, Japan Anti-Tuberculosis Association, Tokyo, Japan

**Keywords:** MAFB, *Mycobacterium tuberculosis*, conditional knockout mouse, mRNA sequencing, host defense

## Abstract

v-Maf avian musculoaponeurotic fibrosarcoma oncogene homolog B (*MAFB*) is a candidate gene associated with early tuberculosis onset identified by a genome-wide association study. Here, we investigated the role of *Mafb* in susceptibility to *Mycobacterium tuberculosis* (*Mtb*) infection in myeloid-specific *Mafb*-knockout (*Mafb*-cKO) mice. We infected bone marrow-derived macrophages (BMMs) from *Mafb*-cKO mice and *Mafb*-cKO mice with *Mtb*. The absence of *Mafb* promoted *Mtb* proliferation in BMMs. RNA sequencing (RNA-seq) revealed activation of the metabolic process and impairment of the response to type Ⅰ interferons (IFNs) in *Mtb*-infected BMMs from *Mafb*-cKO mice, which conforms to our previous findings in *Mtb*-infected human macrophages with *MAFB* knockdown. *Mafb* deficiency increased mortality and bacterial burden in the lungs and spleens during *Mtb* infection in mice. RNA-seq revealed weakened leukocyte or lymphocyte chemotaxis in *Mtb*-infected *Mafb*-cKO mouse lungs. Flow cytometry demonstrated an alteration in the proportion of immune cells in *Mtb*-infected mouse lungs due to *Mafb* deficiency. Together, *Mafb* in myeloid cells is involved not only in the functional antibacterial process of macrophages but also in immune cell recruitment in the lungs, thereby contributing to host defense against *Mtb* infection.

## Introduction

Tuberculosis (TB), caused by *Mycobacterium tuberculosis* (*Mtb*) infection, has resurged as the leading infectious disease, with 8.2 million newly diagnosed cases and 1.2 million deaths in 2023 alone (1). The newly proposed spectrum of TB categorizes this disease into five stages based on the level of infectiousness, presenting symptoms, and macroscopic pathology (2). In this category, subclinical TB is accountable for half the TB burden in any community, making it potentially infectious without any symptoms (3). Therefore, early diagnosis, preventive therapy, or treatment of subclinical TB is critical to prevent further transmission and to ultimately achieve global TB elimination (4). Several studies have attempted to estimate the activation risk based on gene signatures or transcriptional biomarkers (5). Notably, it is important to investigate the factors that determine the trajectory of infection in TB patients or drive susceptibility to TB so as to understand this complex disease as well as to accelerate research in drug and vaccine development.

To date, numerous genome-wide association studies (GWASs) have been conducted to investigate the host genetic factors in TB susceptibility. However, only a few associations have proven reproducibility owing to the modest population sizes, variability in phenotyping across studies, population-specific effects, or complex population structures under certain high-burden settings (6). A meta-analysis combining two GWASs in Thai and Japanese populations did not replicate the association of 25 selected single-nucleotide polymorphisms (SNPs) (7). However, the age-stratified analysis from the same dataset revealed a significant locus on chromosome 20q12 linked to the younger onset group. This locus is located approximately 450-kb upstream of *v-maf avian musculoaponeurotic fibrosarcoma oncogene homolog B* (*MAFB*). Early-onset of TB implies the relatively sooner development after exposure to *Mtb*. The GWAS result suggests that *MAFB* plays a role in the host immunity toward controlling *Mtb* infection. With this background, we investigated the role of *MAFB* as a promising candidate gene involved in TB susceptibility.

MAFB belongs to the large Maf family of transcription factors characterized by a conserved basic leucine zipper (bZip) enabling specific DNA binding to Maf-recognition elements (MAREs) (8). *Mafb* was first identified as a causative gene for segmentation abnormalities in the hindbrain and in defective inner ear development (9). Lethal phenotypes observed in *Mafb*-null mice, including respiratory failure, renal dysgenesis, and parathyroid abnormalities, indicate the critical role of *Mafb* in organogenesis (10). *Mafb* has been demonstrated to control the expression of apoptosis inhibitor of macrophages (AIM) and promote atherosclerosis through inhibiting foam-cell apoptosis (11). In addition, *Mafb* plays diverse roles in various biological processes (BPs), including the myeloid commitment of hematopoietic stem cells, self-renewal of macrophages, generation of thymus, or the creation of hair cuticles (10).

In the context of immune regulation and infectious disease, *MAFB* has been reported to control antiviral response and macrophage polarization (12, 13). Previously, we investigated the function of *MAFB* in *Mtb*-infected human macrophages to explore the biological mechanism underlying *MAFB* in macrophages (14). Our gene knockdown (KD) experiments revealed that MAFB regulates the gene expression related to interferon (IFN) responses in *Mtb*-infected macrophages. In the present study, we investigated the role of *MAFB*, particularly in disease outcomes and dynamic immune cell interactions in organisms by using myeloid-specific *Mafb*-knockout (*Mafb*-cKO) mice (15). We monitored the survival and bacterial burden in the murine organs and found that *Mafb*-cKO mice had higher mortality and bacterial burden during the *Mtb* infection. RNA sequencing (RNA-seq) of *Mtb*-infected *Mafb*-cKO mouse lungs revealed a disrupted chemotaxis. These results were consistent with altered immune cell populations in the lungs of *Mtb*-infected *Mafb*-cKO mice. Taken together, this study highlights *MAFB* as an important gene in macrophages that contributes to protective immunity against *Mtb* infection.

## Materials and Methods

### Ethics statement

Animal experiments in this study were approved by the Animal Care and Use Committee of the Research Institute of Tuberculosis (RIT) (permit number: No. 2021-04). Animals were treated in accordance with the ethical guidelines of RIT. The endpoints were set to determine whether the mice were imminently dying of *Mtb* infection and/or required compassionate euthanasia: bodyweight loss >20% of the initial bodyweight at the time of infection.

### Mice

Macrophage-specific *Mafb* conditional-knockout (*Mafb*^f/f^::LysM-Cre^+/+^ or *Mafb*^f/f^::LysM-Cre^+/-^, *Mafb*-cKO) and *Mafb*^f/f^ control mice were used (15). *Mafb*-cKO and control mice (*Mafb^f/f^*) were maintained in a filtered-air laminar-flow cabinet and provided with sterile bedding, water, and mouse chow at an RIT animal facility. Wild-type (WT) C57BL/6J mice were obtained from The Jackson Laboratory Japan, Inc.

Specific pathogen-free status was verified by testing sentinel mice housed within the colony.

### *Mtb* culture

The *Mtb* strain Erdman was used and stored as previously described (16–18). For determining the bacterial burden in macrophages, the infected cells were lysed with PBS containing 0.1% SDS. Infected lungs or spleens were homogenized using a ShakeMaster Neo (Bio Medical Science). The resulting cell lysates or homogenates were serially diluted and plated in duplicate on 7H10 or 7H11 agar plates supplemented with 10% Middlebrook OADC (BD Bioscience) and 0.5% glycerol. *Mtb* colony-forming units (CFUs) were determined by calculating the mean CFU count from the two plates at each dilution.

### Bone marrow-derived macrophage (BMM) isolation

BMMs were differentiated as described previously (19), with some modifications. Briefly, bone marrow was isolated from the hind legs of *Mafb*-cKO and control mice (6 weeks), washed, and suspended into a single cell by passing through a cell strainer. The bone marrow cells were then incubated at a concentration of 2 × 10^6^ cells/mL in DMEM (Sigma-Aldrich) supplemented with 10% inactivated-fetal bovine serum (FBS, JRH Biosciences Inc.) and 10% of L929-conditioned medium in a 12-well plate for 7 days. Differentiated macrophages in DMEM containing 10% FBS were infected with *Mtb* at a multiplicity of infection (MOI) of one. At one day postinfection (p.i.), BMMs were collected for mRNA sequencing (mRNA-seq). At 1, 3, and 7 days p.i., the number of the intracellular bacteria within BMMs was determined by CFU.

### *Mtb* infection in mice

The experimental mice (age: 6–10 weeks) were transferred to a biosafety level 3 animal facility at RIT. The mice were infected with *Mtb* via the aerosol route using an inhalation exposure system (Glas-Col). This method routinely gave *Mtb* infection at 100–200 CFU per lung one day p.i.

### Survival study

WT and *Mafb*-cKO mice infected with *Mtb* were monitored for 315 days. The mice that survived throughout the experiments or met the endpoint were euthanized by exsanguination under anesthesia with 0.75 mg/kg of medetomidine, 4.0 mg/kg of midazolam, and 5.0 mg/kg of butorphanol *via* the intraperitoneal route (20). Survival probabilities between the two groups were analyzed using Kaplan–Meier analysis and the log-rank test.

### mRNA-seq

mRNA-seq was performed as described previously (18). Briefly, infected BMMs or lungs were homogenized with TRIzol Reagent (Invitrogen), followed by RNA purification using an RNeasy Mini kit (Qiagen). Total RNA qualified and quantified by a 4150 TapeStation system (Agilent) and a Qubit 4 Fluorometer (Invitrogen), respectively, was subjected to construct cDNA libraries using NEBNext® Poly(A) mRNA Magnetic Isolation Module (New England Biolabs) and NEB Next Ultra™ II DNA Library Prep Kit for Illumina (New England Biolabs). All the cDNA libraries were examined for quality using a 4150 TapeStation system and quantified with a Qubit 4 Fluorometer. The libraries were sequenced with a NextSeq1000 system (Illumina).

### Data processing

For read-quality trimming, raw reads were processed with Trim Galore v0.6.7 (https://github.com/FelixKrueger/TrimGalore). The expressions of transcripts were estimated by Salmon v0.12.0 directly from the processed reads (21). Transcript-level expression data was then summarized into gene-level data by the R package tximport v1.30.0 (https://github.com/thelovelab/tximport) in R v4.3.3(22). The analysis for differentially expressed genes (DEGs) was performed by edgeR v4.0.16 (23) using generalized linear models and quasi-likelihood tests (24). The DEGs were further utilized to perform Gene Ontology (GO) enrichment analysis to identify enriched BPs using the R package clusterProfiler v4.10 (25). To reduce redundancy among the identified GO BP categories, a simplification method in clusterProfiler was used. The gene concept network of the top 3 upregulated and downregulated GOBP categories was visualized by cnetplot in clusterProfiler. KEGG pathway enrichment analysis was conducted using ShinyGO v0.82 (26), an online gene set enrichment tool (http://bioinformatics.sdstate.edu/go/). Adapted KEGG pathway diagrams were visualized using Pathview v1.42.0 in R software (27). Pathway source: KEGG (https://www.kegg.jp). Gene set enrichment analysis (GSEA) was performed locally using the GSEA desktop application (Broad Institute, v4.2.3) with the WikiPathway gene sets (c2.cp.wikipathways.v2024.1.Hs.symbols.gmt) obtained from the Molecular Signature Database (MSigDB). MafB ChIP-seq peaks (GEO GSM1964739/SRA SRX1465586) (28) were downloaded via ChIP-Atlas (accessed 27 June 2025) (29) and compared with the DEGs identified in the present study. The ChIP-seq Atlas is accessible at https://chip-atlas.org/.

### Flow cytometry

Infected lung cells were obtained using the Lung Dissociation Kit (Miltenyi Biotec) according to the manufacturer’s instructions. Briefly, cell suspensions from the lungs were incubated with ACK buffer to lyse red blood cells. The cells were washed and diluted in MACS buffer (PBS supplemented with 2mM EDTA and 2% FBS) to achieve 1–3 × 10^6^ cells/mL. The cells were incubated with TruStain FcX™ PLUS (anti-mouse CD16/32) (Biolegend), followed by staining with antibodies against CD4, CD8, CD45R, CD3, SiglecF, CD64, CD11b, CD45, or Ly6G (BioLegend). The stained cells were then fixed with the fixation buffer (BioLegend) to inactivate infected *Mtb* for 24 h at 4°C. The cells were analyzed on a BD FACSLyric^TM^ using analysis software BD FACSuite™ Application V1.4.0.7047 and FlowJo™ Software v10.10 (BD Biosciences).

## Results

### *Mafb* deficiency on mycobacterial killing in macrophages

In our previous study, we demonstrated impaired inflammatory responses in *MAFB* knockdown, PMA-stimulated THP-1 cells (*MAFB*-KD macrophages). However, no significant difference in bacterial burden was observed between *MAFB*-KD macrophages and control macrophages at 24 h or 48 h p.i., suggesting that the knockdown effect and/or the duration of infection was insufficient to detect intracellular bacterial proliferation (14). In this study, we investigated the effect of *Mafb* deficiency on mycobacterial proliferation in BMMs (**Figure 1**). For this purpose, we infected BMMs of *Mafb*^f/f^::LysM-Cre^+/-^ (*Mafb*-cKO) or *Mafb*^f/f^ (control) with *Mtb* at an MOI of one for 1, 3, and 7 days. At 3 and 7 days p.i., *Mafb*-cKO BMMs exhibited a higher bacterial burden compared to control BMMs. Notably, the intracellular number of *Mtb* in *Mafb*-cKO BMMs increased significantly more rapidly from day 3 to day 7, suggesting that the absence of *Mafb* transforms macrophages into a more permissive environment for *Mtb* proliferation.

**Figure 1.**
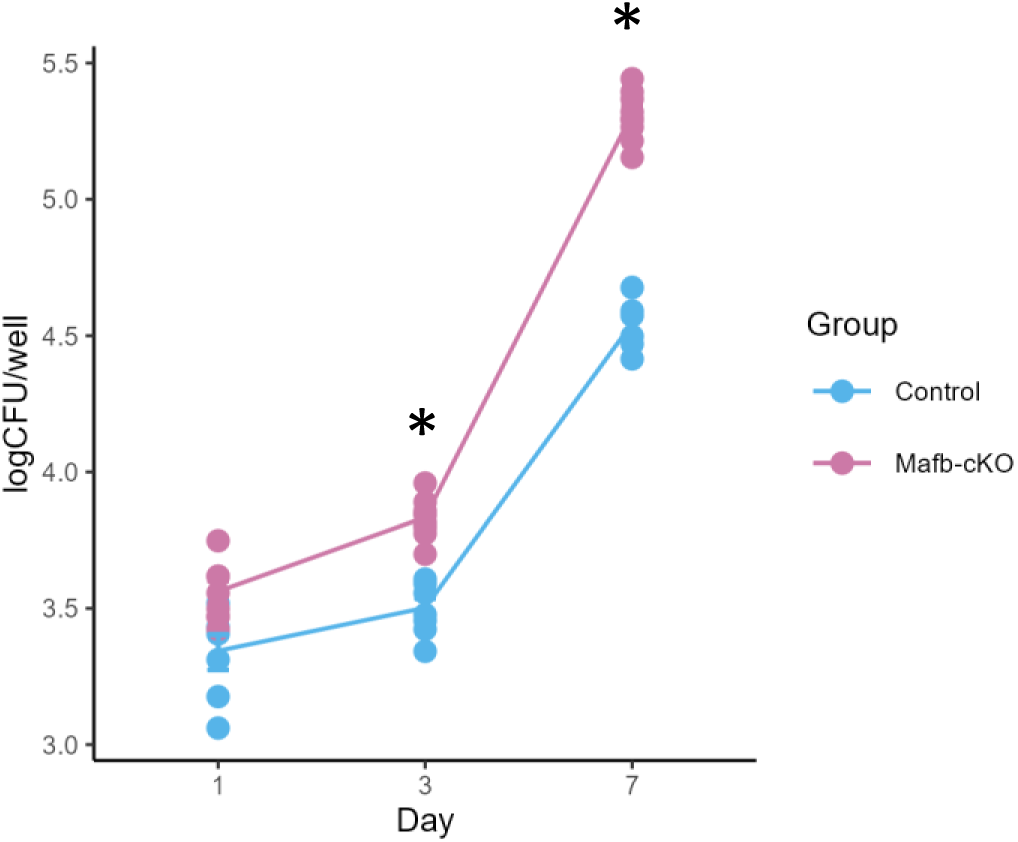
Mycobacterial proliferation on *Mafb*-deficient macrophages. *Mafb*^f/f^::LysM-Cre^+/-^ (*Mafb*-cKO) and *Mafb* ^f/f^ (control) bone marrow-derived macrophages (BMMs) were infected with *Mycobacterium tuberculosis* (*Mtb*). At 1, 3, and 7 days postinfection (p.i.), the numbers of the intracellular bacteria were determined by colony forming units (CFUs). \**P* < 0.01 using Welch’s t-test, with Holm–Bonferroni correction applied for multiple comparisons.

### *Mafb*-cKO BMMs demonstrated functional changes in metabolic process and immune response during *Mtb* infection

When macrophages are exposed to *Mtb*, they internalize the bacteria, and *Mtb* begins adapting to the intracellular environment by 24 h p.i. During this period, macrophages undergo robust transcriptional changes, indicating active host–pathogen interactions (30). To investigate the transcriptional function of *Mafb* in *Mtb*-infected macrophages, we infected BMMs from *Mafb*^f/f^::LysM-Cre^+/-^ (*Mafb*-cKO) and *Mafb*^f/f^ (control) with *Mtb* and conducted mRNA-seq at 24 h p.i. mRNA-seq comparing between *Mtb*-infected BMMs from *Mafb*-cKO and control mice identified 1223 DEGs (**Figure 2A**). GO analysis for BP (GOBP) identified 974 significantly enriched GOBP terms in 614 upregulated DEGs in *Mtb*-infected *Mafb*-cKO BMMs, including leukocyte cell–cell adhesion, reactive oxygen species (ROS) metabolic process, or nucleotide metabolic process (**Figure 2B**). In 609 downregulated DEGs, 493 significantly enriched GOBP terms were identified, including response to virus, defense response to symbiont, or regulation of innate immune response (**Figure 2B**). To visualize the interactions of the genes annotated to each GO term, we constructed gene networks (**Figure 2C**).Upregulated genes annotated to leukocyte cell–cell adhesion include *Itgb2*, *Itgb7*, *Itgal*, *F11r*, *Selplg*, *Cd44* (adhesion molecules or integrins), *Ccr2*, *Cx3cr1*, *Cd86*, *Cd74*, *Cd24a*, *Cd177*, *Cd300a* (leukocyte surface receptors), *H2-Ab1*, *H2-Aa*, *H2-Eb1*, *H2-DMa*, *H2-DMb1* (MHC molecules), and *Arg2*, *Thbs1*, *Alox5*, *Sirpb1b*, *Sirpb1c* (immune modulation), suggesting that the macrophages are in the state where they are actively participating in immune surveillance, cellular communication, and antigen presentation. Genes annotated to ROS metabolic process include *Cybb*, *Cyba*, *Ncf2* (ROS generation), *Prdx1*, *Prdx5*, *Prdx6*, *Txn1*, *Txnrd1*, *Sod2*, *G6pdx*, *Gpx3*, *Nnt* (ROS detoxification, antioxidant defense), and *Thbs1*, *Rhoa*, *Atg5* (oxidative stress modulation). The downregulated genes annotated to top significantly enriched GOBP terms were highly overlapped: *Apobec1*, *Adar*, *Ifi204*, *Ifi203*, *Ifi205*, *Ifi206*, *Ifi207*, *Ifi208*, *Ifi209*, *Ifi211* (RNA editing and modification), *Ifih1*, *Dhx58*, *Rigi*, *Ddx60*, *Stat1*, *Stat2*, *Irf3*, *Irf7*, *Tbk1*, *Trim25*, *Trim5*, *Trim21*, *Trim30a*, *Trim30c*, *Trim30d*, *Trim34a*, *Trim35*, *Trim12c*, *Trim8*, *Isg15*, *Isg20*, *Oas1a*, *Oas1g*, *Oas2*, *Oas3*, *Oasl2*, *Ifit1*, *Ifit2*, *Ifit3*, *Ifit3b*, *Ifit1bl1*, *Mx1*, *Mx2*, *Bst2*, *Dtx3l*, *Rnasel*, *Parp9*, *Shfl*, *Slfn8*, *Slfn9*, *Gbp3*, *Gbp5*, *Gbp7*, *Rnf135*, *Nt5c3* (innate immune sensors and IFN-stimulated genes (ISGs)), *Irgm1*, *Irgm2*, *Igtp*, *Mov10* (autophagy and intracellular pathogen defense), *Pou2f2*, *Il10rb*, *Il15*, *Tlr9*, *Tlr1*, *Cd37*, *Cd180*, *Cd36*, *Cd300ld3*, *Pik3ap1*, *Rtp4*, *Sash1*, *Trafd1*, *Appl2* (transcription factors and signal transduction), *Eif2ak2*, *Pml*, *Bcl2l1*, *Trex1*, *Xaf1*, *Htra2*, *F830016B08Rik* (cell cycle, apoptosis, and DNA repair), *Lacc1*, *Apoe*, *Adam8*, and *Hyal1* (metabolism and miscellaneous functions). These downregulated DEGs suggested weakened pathogen sensing and reduced IFN response or inflammatory signaling, indicating a potential shift to an anti-inflammatory phenotype in *Mtb*-infected BMMs of *Mafb*-cKO mice. KEGG pathway enrichment analysis revealed that oxidative phosphorylation and chemical carcinogenesis-ROS were enriched in the upregulated DEGs (**Supplementary Figure 1A**), whereas ECM-receptor interaction, lysosome, and endocytosis were enriched in downregulated DEGs (**Supplementary Figure 1B**). As depicted in the lysosome pathway diagram, the proton pump ATPeV, which plays a critical role in lysosomal acidification, is downregulated (**Supplementary Figure 2**) (31). By GSEA using all expressed genes, a pathway of immune response to TB (https://www.wikipathways.org/instance/WP4197) exhibited impairment in *Mtb*-infected *Mafb*-cKO BMMs (**Supplementary Figure 3**). These enriched BPs and pathways are consistent with the previous results obtained from mRNA-seq of *Mtb*-infected *MAFB*-KD macrophages (14). IFN-gamma inducible chemokines (*Cxcl11*, *Ccl2*, *Ccl7*, *Cxcl9*, *Cxcl10*) were downregulated in *Mtb*-infected *MAFB*-KD macrophages, as well as in *Mtb*-infected BMMs from *Mafb*-cKO mice (**Supplementary Figure 4**). Thus, we proposed that MAFB in human and murine macrophages has a similar role during *Mtb* infection.

**Figure 2.**
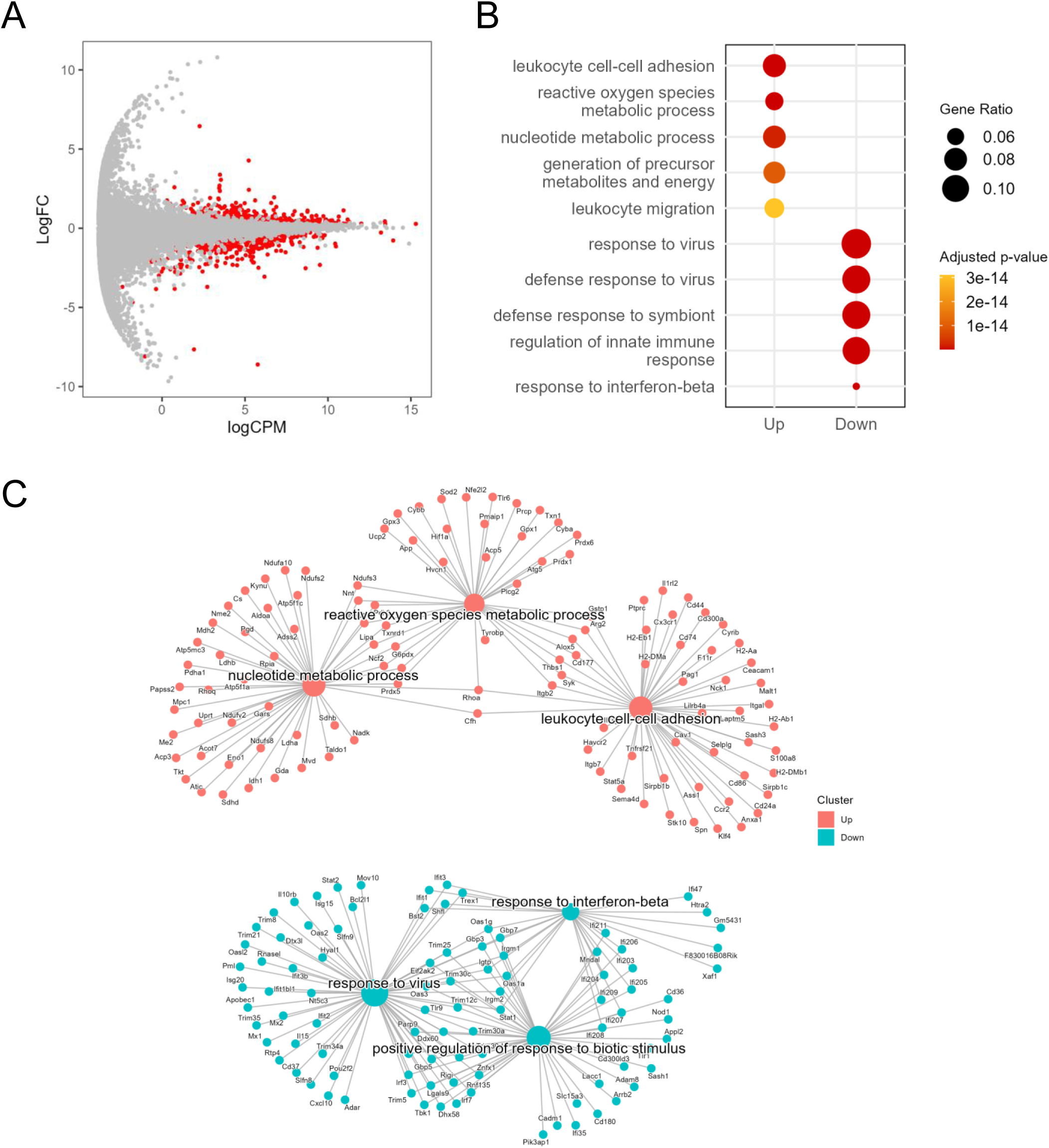
Transcriptomics of *Mtb*-infected *Mafb*-cKO bone marrow-derived macrophages (BMMs) mRNA sequencing (mRNA-seq) of *Mtb*-infected *Mafb*-cKO BMMs was performed.MA plot showing 1223 differentially expressed genes (DEGs) in *Mtb*-infected *Mafb*-cKO BMMs compared to those in *Mtb*-infected *Mafb* ^f/f^ control BMMs, marked in red (FDR < 0.01). Each dot represents expressed genes in the sample. Log FC, log fold change. LogCPM, log count per million. (B) Gene Ontology (GO) analysis for upregulated or downregulated DEGs. Enriched GO biological process (BP) categories in *Mtb*-infected *Mafb*-cKO BMMs can be seen. (C) Gene concept networks of the top 3 upregulated (Up) and downregulated (Down) GOBP categories in *Mtb*-infected *Mafb*-cKO BMMs. Upregulated GOBP categories are leukocyte cell–cell adhesion, reactive oxygen species metabolic process, and nucleotide metabolic process, colored in salmon pink. Downregulated GOBP categories are response to virus, defense response to symbiont, and regulation of innate immune response, as shown in blue.

### *Mafb* deficiency in macrophages increased mortality during *Mtb* infection in mice

To examine whether *Mafb* deficiency in macrophages influences the outcome of *Mtb* infection in mice, we conducted aerosol infections in *Mafb*-cKO mice and WT mice and monitored them for 315 days. We found that *Mafb*-cKO mice started succumbing after 20 weeks, and no *Mafb*-cKO mice survived by the end of the experiment (**Figure 3A**). The survival probability between the two groups was analyzed by Kaplan–Meier analysis and the log-rank test. The median survival of *Mafb*-cKO mice was 211 days, which was significantly shorter than that of WT mice. We next determined the bacterial burden in the murine organs after *Mtb* infection. At 10 and 20 weeks p.i., *Mafb*-cKO mice exhibited significantly higher burden in the lungs, but not at 4 weeks p.i. (**Figure 3B**). The spleens of *Mafb*-cKO mice also showed a higher burden at 10 and 20 weeks p.i., demonstrating the involvement of *Mafb* in the control of the bacterial burden in the lung and spleen (**Supplementary Figure 5**).

**Figure 3.**
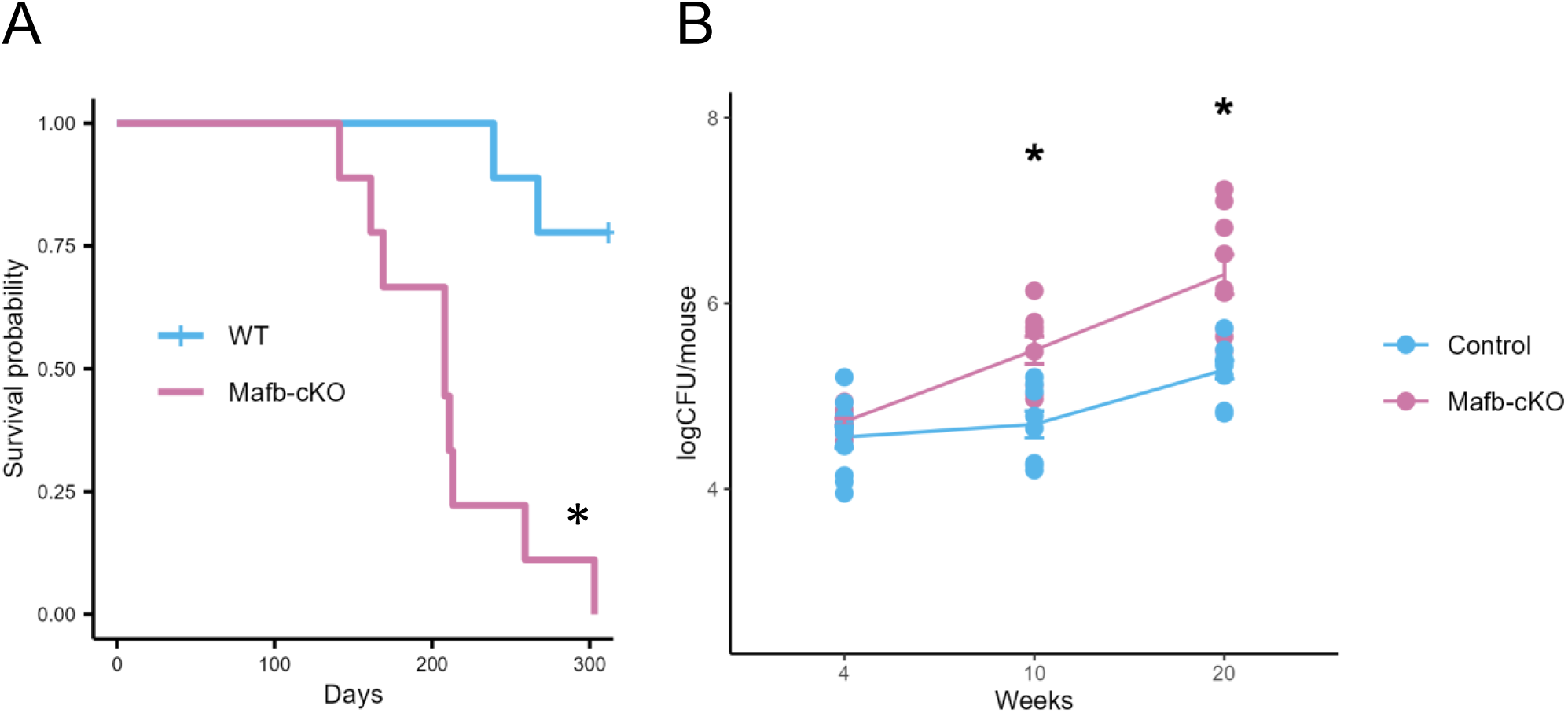
Effect of macrophage-specific *Mafb* deficiency on TB susceptibility in mice. (A) *Mafb*^f/f^::LysM-Cre^+/+^ (*Mafb*-cKO) mice and wild-type (WT) mice (*n*=9/group) were aerosol-infected with *Mtb*, and their survival was monitored for 315 days. Survival probability between the two groups was analyzed by Kaplan–Meier analysis and the log-rank test. The median survival of *Mafb*-cKO mice was 211 days. Most of the WT mice survived when the experiment was ended. \**P* = 0.00011. (B) Bacterial loads in the lungs of *Mtb*-infected *Mafb*^f/f^::LysM-Cre^+/-^ (*Mafb*-cKO) and control mice were determined by CFU at 4, 10, and 20 weeks p.i. Data from individual mice is shown. \**P* < 0.01 using Tukey–Kramer test.

### Transcriptomics of *Mtb*-infected *Mafb*-cKO mouse lungs

To investigate whether *Mafb* deficiency in macrophages alters BPs in the lungs during *Mtb* infection, we performed mRNA-seq on the lungs of *Mtb-*infected *Mafb*-cKO and control mice at 10 or 20 weeks p.i., respectively. At 10 weeks p.i., 89 genes were identified as DEGs in the lungs of *Mtb*-infected *Mafb*-cKO mice (**Figure 4A**). Among these 89 genes, 48 genes were upregulated and 41 genes were downregulated. GOBP of DEGs demonstrated that cell–cell adhesion, leukocyte proliferation, or the regulation of T-cell activation were activated, whereas complement activation, cellular response to type Ⅱ IFN, synapse pruning, and response to protozoan were suppressed in the lungs of *Mafb*-cKO mice (**Figure 4B**). Concept gene network for GO categories visualized that *Cd1d1*, *Cdkn2a*, *Tarm1*, *Havcr2*, and *Slfn1* were the key genes for T-cell regulation (**Figure 4C**). KEGG pathway enrichment analysis demonstrated the enrichment of osteoclast differentiation (**Supplementary Figure 6A**). Complement components such as *C1qa*, *C1qb*, or *C1qc*, identified in suppressed GO categories, played a central role in complement activation. The involvement of *Mafb* in regulating complement components was consistent with the previous report (15). In addition to complement components, downregulated DEGs included cytokine ligands such as *Ccl8*, which is also known as *monocyte chemoattractant protein 2* (MCP2), *Ccl12*, known as *monocyte chemoattractant protein 5* (MCP5), or *Pf4,* known as *Cxcl4*. KEGG pathway enrichment analysis demonstrated that complement and coagulation cascade, and chemokine signaling pathway were enriched in downregulated DEGs (**Supplementary Figure 6BC**). At 20 weeks p.i., 267 DEGs were identified (**Figure 5A**), of which DEGs found at 10 weeks p.i. were included. Among the 267 genes, 110 genes were upregulated and 157 genes were downregulated. GOBP showed that myeloid leukocyte activation, myeloid leukocyte differentiation, the regulation of macrophage activation, the regulation of endocytosis, and the regulation of angiogenesis were upregulated, whereas leukocyte migration, leukocyte chemotaxis, leukocyte proliferation, and immune response cell-surface receptor-signaling pathway were downregulated in the lungs of *Mtb*-infected *Mafb-*cKO mice (**Figure 5B**). Concept gene network revealed *Csfs* (GM-CSF), a key regulator for macrophage and dendritic cell function, *Mmp8*, *Cd177*, genes associated with neutrophil activation and migration, or *Sirpb1 family* for phagocytosis and immune modulation in upregulated DEGs, highlights strong differentiation and the activation of myeloid-derived immune cells (**Figure 5C**). Down regulated DEGs included *Ccl22*, *Ccl8*, *Ccl5*, *Cx3cr1*, *Pf4*, and *Ccr7*, which are involved in chemokine signaling or leukocyte migration; *P2rx7*, *Nfatc2*, and *Ptpn22*, regulatory genes in T-cell activation and immune tolerance, *Cd22* or *Icosl*, which are involved in B cell-mediated immune response, suggesting reduced adaptive immune activation and leukocyte or lymphocyte recruitment in the lungs of *Mtb*-infected *Mafb-*cKO mice compared to those of control mice (**Figure 5C**).

**Figure 4.**
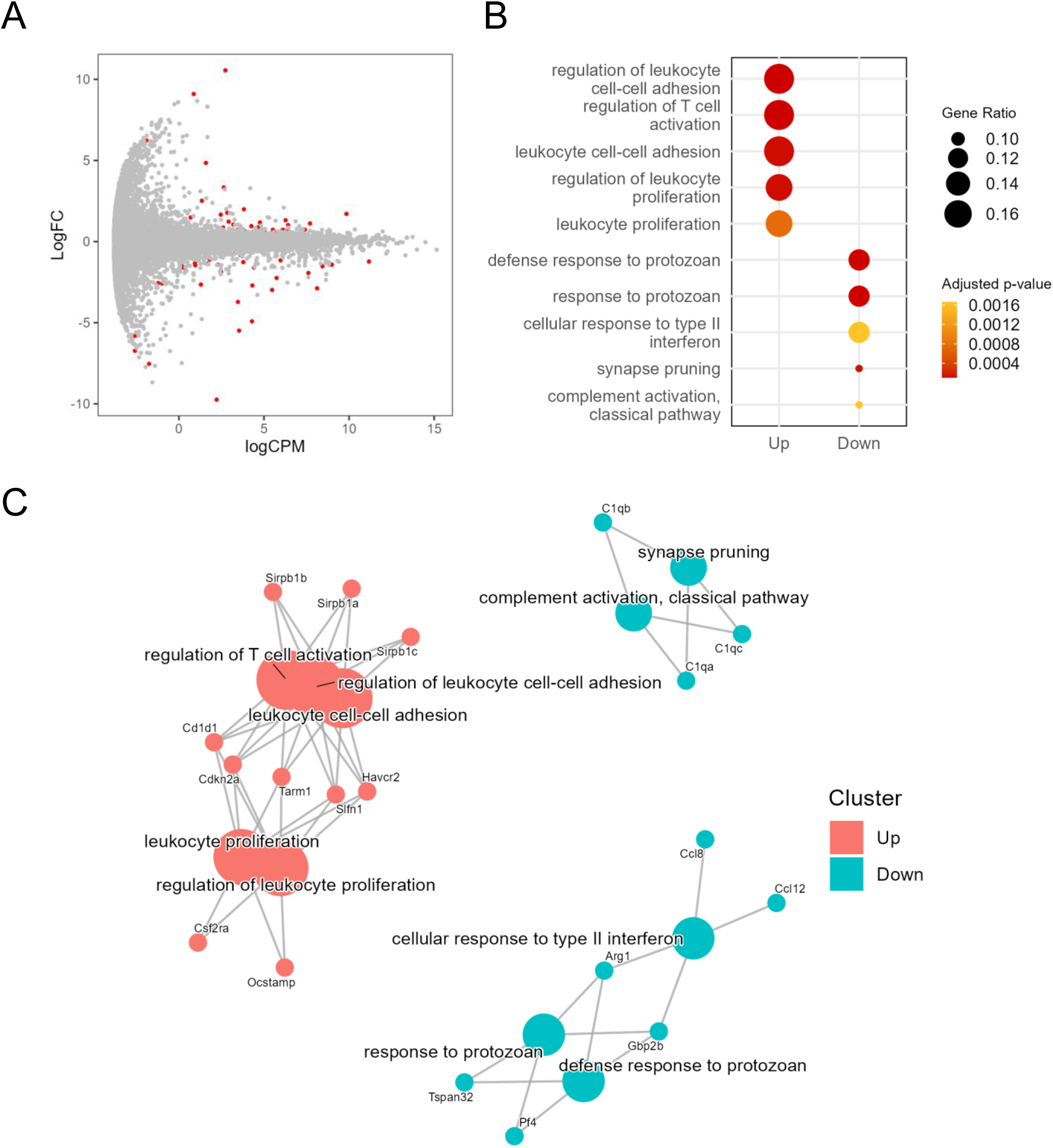
Transcriptomics of *Mtb*-infected *Mafb*-cKO mouse lung at 10 weeks p.i. *Mafb*-cKO mice and control mice were aerosol-infected with *Mtb* for 10 weeks. (A) MA plot showing 89 DEGs in *Mtb*-infected *Mafb*-cKO mouse lungs compared to those in *Mtb*-infected control mouse lungs, marked in red (FDR < 0.01). Each dot represents expressed genes in the sample. Log FC, log fold change. LogCPM, log count per million. (B) GO analysis for DEGs. Enriched GOBP categories in *Mtb*-infected *Mafb*-cKO lungs are shown. (C) Gene concept network of the top 3 upregulated (Up) and downregulated (Down) GOBP categories in *Mtb*-infected *Mafb*-cKO mouse lungs. Upregulated GOBP categories include regulation of leukocyte cell–cell adhesion, regulation of T-cell activation, leukocyte cell–cell adhesion, and leukocyte proliferation, colored in salmon pink. Downregulated GOBP categories are defense response to protozoan, response to protozoan, cellular response to type Ⅱ interferon, synapse pruning, and complement activation, and classical pathway, as shown in blue.

**Figure 5.**
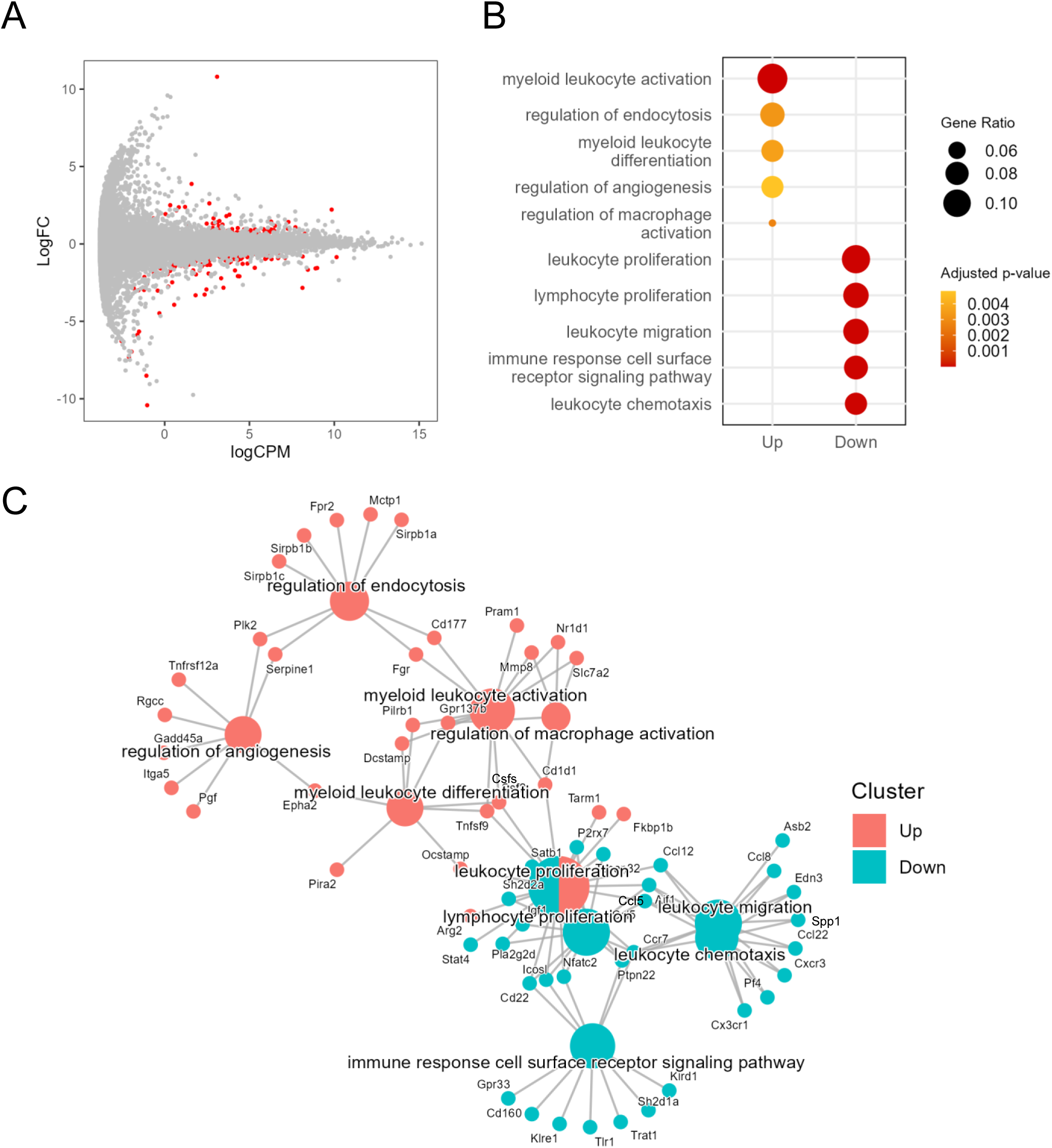
Transcriptomics of *Mtb*-infected *Mafb*-cKO mouse lung at 20 weeks p.i. *Mafb*-cKO mice and control mice were infected with an aerosol of *Mtb* for 20 weeks. (A) MA plot showing 267 DEGs in *Mtb*-infected *Mafb*-cKO mouse lungs in comparison with control mouse lungs marked in red (FDR < 0.01). Each dot represents expressed genes in the sample. Log FC, log fold change. LogCPM, log count per million. (B) GO analysis for DEGs. Enriched GOBP categories in *Mtb*-infected *Mafb*-cKO lungs were shown. (C) Gene concept network of the top 3 upregulated (Up) and downregulated (Down) GOBP categories in *Mtb*-infected *Mafb*-cKO mouse lungs. The upregulated GOBP categories include myeloid leukocyte activation, regulation of endocytosis, myeloid leukocyte differentiation, regulation of angiogenesis, and regulation of macrophage activation colored in salmon pink. Downregulated GOBP categories are leukocyte proliferation, lymphocyte proliferation, immune response cell surface receptor signaling pathway, and leukocyte chemotaxis, shown in blue.

### Immune cell recruitment in the lungs of *Mtb*-infected *Mafb*-cKO mice

Transcriptomics of the lungs of *Mtb*-infected *Mafb*-cKO mice suggested altered recruitment of immune cells during *Mtb* infection (**Figure 4 and 5**). Therefore, we investigated the proportion of immune cells in the lungs of *Mafb*-cKO mice during *Mtb* infection by flow cytometry (**Figure 6**). The frequencies of both CD4^+^ and CD8^+^ T-cells were high at 10 weeks p.i., and then decreased at 20 weeks p.i. in control mice, whereas they were at the same levels in *Mafb*-cKO mice during infection, suggesting blocked early recruitment of CD4^+^ and CD8^+^ T-cells in the infected lungs of *Mafb*-cKO mice in line with the transcriptomic feature of negative T cell regulation at 10 weeks p.i. (**Figure 6**). The frequency of B-cells remained the same from 10 weeks to 20 weeks p.i. in the control mice; however, it decreased in *Mafb*-cKO mice at 20 weeks p.i., which also supports the transcriptomics data. Despite the impaired chemokine signaling, the frequency of neutrophils was significantly higher in *Mafb*-cKO at 10 or 20 weeks p.i. compared to that in control mice. Although the frequency of macrophages was slightly lower in *Mafb*-cKO mice, the difference was not statistically significant due to the high inter-sample variability. Nonetheless, these results indicate that *Mafb* deficiency in macrophages affects the recruitment of various immune cells to the lungs of *Mtb*-infected mice.

**Figure 6.**
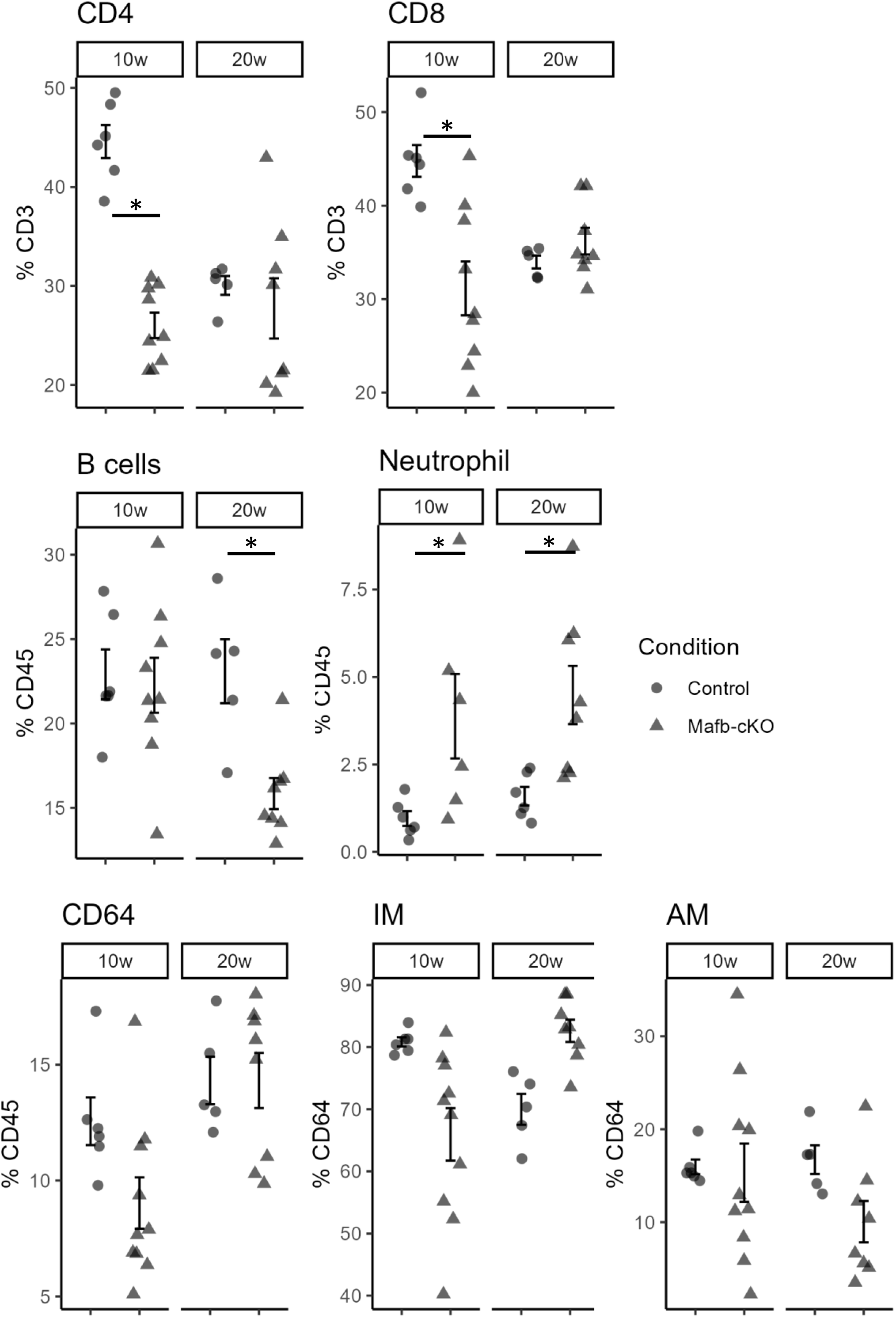
Population of immune cells in *Mtb*-infected Mafb-cKO mice. The proportion of the immune cell population in the lungs of *Mtb-*infected *Maf*b-cKO or control mice was determined by flow cytometry at 10 or 20 weeks p.i. The proportions of CD45^+^, B-cells, neutrophils, macrophages, CD4^+^ T-cells, CD8^+^ T-cells, and alveolar macrophages are shown. \**P* < 0.05 using the Wilcox test for each time point.

## Discussion

We have demonstrated that *MAFB* deficiency activates oxidative phosphorylation and impairs type Ⅰ and Ⅱ IFN responses in *Mtb*-infected human macrophages (14). For a deeper understanding of *MAFB*’s role in *Mtb* infection, we investigated its function using a murine model of myeloid-specific *Mafb* conditional-knockout (*Mafb*-cKO) mice. Transcriptomics of BMMs from *Mafb*-cKO mice revealed that a ROS metabolic process and oxidative phosphorylation were activated, whereas the IFN responses were suppressed during *Mtb* infection (**Figure 2**), which is consistent with our previous results (14). Therefore, we assume that *MAFB* in human and murine macrophages acts similarly in response to *Mtb* infection.

We found that BMMs from *Mafb*-cKO mice failed to control intracellular *Mtb* proliferation (**Figure 1**). In fact, *Mtb* infection typically induces the production of ROS in infected macrophages to reduce the intracellular bacterial load (32). However, an imbalance between ROS and antioxidants leads to oxidative stress, which contributes to the onset and progression of pulmonary diseases, including TB (33, 34). In BMMs from *Mafb*-cKO mice, the ROS metabolic process was activated, and its related genes were identified. CYBB, CAMK2B, and ITPR1 were involved in ROS production, and SOD2, GPX3, and CAT were involved in ROS clearance (35). CYBB is the major catalytic subunit of nicotinamide adenine dinucleotide phosphate (NADPH) oxidase, encoding NOX2 that possesses antimicrobial activity against *Mtb* (36). The superoxide dismutase expressed by *Sod2* detoxifies the major ROS to protect host cells from the damage caused by excessive ROS. Paradoxically, the overexpression of *Sod2* promotes the intracellular survival of *Mtb* (37). During *Mtb* infection, genes related to both ROS production and clearance were upregulated in BMMs derived from *Mafb*-cKO mice, implying that BMMs from *Mafb*-cKO mice generate ROS to clear pathogen while maintaining redox balance to prevent self-damage during infection, thereby actively attempting to eliminate the excessive ROS.

*Mtb* infection also activates antiviral responses, including the induction of type I IFNs, in infected macrophages (38, 39). During infection, mycobacterial DNA is initially released from the phagosomes into the cytosols, where it is recognized by cyclic GMP-AMP synthase (cGAS), initiating type I IFN production. This recognition triggers the activation of the cGAS-STING-TBK1 cascade and transcription factors IRF3 and IRF7, followed by the production of type Ⅰ IFNs and other cytokines (40).

Activated IRF3 translocates into the nucleus and binds to IFN-stimulated response element (ISRE) in the promoters of type Ⅰ IFNs and proinflammatory genes for further transcriptional induction (41). It has been shown that IRF3 is essential for downstream genes, such as *Cxcl10* and *Ifit1*, which are induced by IFN-β and IFN-γ (42). In *Mtb*-infected BMMs from *Mafb*-cKO mice, *Tbk1*, *Irf3*, *Irf7*, *Stat1*, *Stat2,* and other genes with ISRE (*Adar*, *Bst2*, *Cxcl10*, *Gbp2*, *Ifit1*, *Ifit2*, *Ifit3*, *Irf7*, *Irf9*, *Isg15*, *Isg20*, *Mx1*, *Mx2*, *Oas2*, *Oas3*, *Rsad2*, *Sp100)* and type I IFNs (*IFNα*, *IFNβ*) were significantly downregulated (**Figure 2**, **Supplementary Figure 7A**), suggesting that *Mafb* can be a positive regulator in the cGAS-STING-TBK1 cascade.

In mycobacterial infection, type I IFNs can be detrimental to the host, considering the type Ⅰ IFN-driven susceptibility in the mouse model carrying the sensitive allele of the *Sst1* locus (43). Contrary to the results of BMMs from *Mafb*-cKO mice, type Ⅰ IFNs (IFNα) or genes with ISRE were found to be upregulated in *Mtb*-infected *Mafb*-cKO mouse lungs **(Supplementary Figure 7BC**). This result may explain the increased mortality and higher bacterial burden in the organs of *Mtb*-infected *Mafb*-cKO mice (**Figure 3**). A possible explanation for this higher bacterial burden in BMMs from *Mafb*-cKO mice is weakened lysosome biogenesis demonstrated by a pathway analysis (**Supplementary Figure 1B**, **2**). In addition, genes related to the immune response to the TB pathway were downregulated in *Mafb*-cKO BMMs compared to those in control BMMs (**Supplementary Figure 3**). Together with the imbalanced ROS metabolism, we proposed that *Mafb*-deficient BMMs lack an effective antibacterial response that would contain intracellular *Mtb*.

As expected from the failure of *Mtb* containment at a cellular level, *Mafb* depletion in macrophages causes increased mortality and a higher bacterial burden in the organs of *Mtb*-infected mice (**Figure 3**). To gain an insight into the mechanism underlying the susceptibility of *Mafb*-cKO mice, we performed RNA-seq of *Mafb*-cKO mouse lungs. We identified 1223 DEGs by comparing between *Mtb*-infected BMMs from *Mafb*-cKO and control mice. In the lungs of *Mtb*-infected *Mafb*-cKO mice, 89 or 267 genes were identified as DEGs at 10 weeks or 20 weeks p.i., respectively. MafB is a transcription factor that binds to Maf recognition elements (MAREs) in gene promoters (8). We obtained MafB ChIP-sequencing data (28) to compare with the DEGs identified in the present study. Among the 1,223 DEGs in *Mtb*-infected *Mafb*-cKO BMMs, 413 genes (33.8%) overlapped with MafB target genes, including 175 upregulated and 238 downregulated genes (**Table 1**). In the lungs of *Mtb*-infected *Mafb*-cKO mice, 28 DEGs (31.5%) at 10 weeks p.i. and 55 DEGs (20.6%) at 20 weeks p.i. overlapped with MafB target genes, respectively (**Table 1**). The proportion of DEGs directly bound by MafB was comparable between *Mafb*-cKO BMMs and lungs at 10 weeks p.i.; however, this proportion was lower at 20 weeks p.i., despite a greater number of DEGs being detected. This suggests that secondary effects of *Mafb* deficiency may contribute to the increased mortality observed in *Mafb*-cKO mice during the later stage of infection.

**Table 1.** Comparison of MafB ChIP-seq peaks and the DEGs. MafB ChIP-seq peaks are compared with the DEGs identified in *Mtb*-infected *Mafb*-cKO mouse BMMs, *Mtb*-infected *Mafb*-cKO mouse lungs at 10 weeks p.i., or *Mtb*-infected *Mafb*-cKO mouse lungs at 20 weeks p.i. The ChIP-seq data (GEO GSM1964739/SRA SRX1465586) (28) was downloaded via ChIP-Atlas (accessed 27 June 2025) (29)

|  |  | Upregulated | Downregulated | Total |
| --- | --- | --- | --- | --- |
| *Mafb*-cKO mouse BMMs | DEGs | 614 | 609 | 1223 |
|  | MafB ChIP-seq peaks | 175 | 238 | 413 |
|  |  | 28.5% | 39.1% | 33.8% |
| *Mafb*-cKO mouse lungs, 10w | DEGs | 48 | 41 | 89 |
|  | MafB ChIP-seq peak | 14 | 14 | 28 |
|  |  | 29.2% | 34.1% | 31.5% |
| *Mafb*-cKO mouse lungs, 20w | DEGs | 110 | 157 | 267 |
|  | MafB ChIP-seq peak | 24 | 31 | 55 |
|  |  | 21.8% | 19.7% | 20.6% |

Transcriptomics of the lungs of *Mtb*-infected *Mafb*-cKO mice displayed activated myeloid-derived immune cells and differentiation (**Figure 4**). This finding is consistent with the reports that low MafB levels activate self-renewal in resident macrophages(28, 44). Vanneste *et al.* exhibited that myeloid-specific *Mafb* deletion increased both the proliferative ability and cell death in macrophages, decreasing number of macrophages in the mouse lungs (45). We also demonstrated a slightly decreased population of CD64^+^ macrophages among immune cells at 10 weeks p.i. but not at 20 weeks p.i. (**Figure 6**). We displayed a significantly higher frequency of neutrophils in *Mafb*-cKO mouse lungs (**Figure 6**). The result is consistent with the necrosis of macrophages and bacillary replication induce neutrophil recruitment (46). In fact, neutrophil accumulation correlates with increased disease severity, suggesting that excessive neutrophils may exacerbate TB pathology (47). *Mtb*-infected *Mafb*-cKO mice displayed a higher bacterial burden at 10 weeks p.i. in the lungs. Neutrophil recruitment is further enhanced by their release of mediators in response to *Mtb* (48). These findings explain that neutrophils can drive further neutrophil recruitment to the lungs of *Mafb*-cKO mice. Kanai *et al.* also demonstrated an increased myeloid-cell infiltration, including neutrophils, in an ischemic acute kidney injury (AKI) model in *Mafb*-cKO mice, suggesting that *Mafb* is involved in myeloid-cell migration both in the site of infection and injury. (49) Considering that *Mafb* regulates thermogenesis in brown adipose tissue in *Mafb*-cKO mice under cold conditions (50), it is suggested that *Mafb* controls various homeostatic functions in macrophages under infections, injuries, or cold conditions.

We demonstrated that *Mafb* deficiency in macrophages impaired the cell signaling for leukocyte migration and the recruitment of CD4^+^ and CD8^+^ T-cells in the lungs of *Mtb*-infected mice, suggesting weakened adaptive immunity at an early stage of infection. Several studies have shown the importance of macrophage activation by IFN-γ produced from CD4^+^ T-cells for protective immunity against *Mtb* in mice (51–53). The depletion of CD4^+^ T-cells leads to increased bacterial loads and increased severity of the infection in *Mtb*-infected C57BL/6 mice (54). In a macaque model, CD4^+^ T-cells display an “innate-like” defense system and serve as master helper cells to recruit other Th-like effector cells, thereby successfully preventing early extrapulmonary *Mtb* dissemination (55).

In summary, the present study provides evidence that *Mafb* depletion in myeloid cells not only impairs macrophage bactericidal activity but also disrupts immune cell recruitment, leading to failed bacterial control and higher mortality in *Mtb*-infected mice.

## Author Contributions

HH, SS, MHijikata and NK designed the project. HH, SS, HN, and SO performed the experiments. MHamada, and ST provided the experimental materials. HH and SS analyzed the data. HH, SS, MHijikata and NK wrote and revised the manuscript. All authors approved the manuscript.

## Acknowledgement

We thank Dr. Masayuki Umemura from Ryukyu University for valuable discussions and expert advice on flow cytometric analysis. We also thank all our colleagues and staffs at The Research Institute of Tuberculosis, Japan Anti-Tuberculosis Association for technical and administrative support.

